# AlignmentFold and AlignmentPartition: Improving the align-then-fold approach for RNA secondary structure prediction

**DOI:** 10.1101/2025.07.23.666478

**Authors:** Abhinav Mittal, David H. Mathews

## Abstract

The combination of free energy minimization with sequence covariation analysis is a popular approach for predicting conserved secondary structure using a fixed alignment of homologous RNA sequences. In this work we revisit this approach by developing the new AlignmentFold and AlignmentPartition programs. AlignmentFold and AlignmentPartition predict consensus minimum free energy structures and base pairing probabilities, respectively. We determined new nearest neighbor thermodynamic parameters for gaps and non-canonical base pairs that align with base pairs. These frequently occur for structure prediction with a fixed input alignment. We assessed the impact of alignment prediction method and size on the prediction accuracy. We also assessed the improvement in prediction accuracy conferred by sequence covariation analysis. In contrast to previous solutions, we do not include covariation in the scoring of structure quality and achieve prediction accuracy as good as other tools. AlignmentFold and AlignmentPartition are freely available as part of the RNAstructure software package at https://rna.urmc.rochester.edu/RNAstructure.html.

## INTRODUCTION

RNA sequences play diverse roles in cellular processes ranging from gene expression to anti-viral defense (Cech and Steitz 2014). Consequently, they emerged as drug targets, drugs, and vaccines (Damase et al. 2021). While new classes of non-coding RNAs (ncRNAs) with their functions continue to come to light (Flynn et al. 2021), a better understanding of RNA structure and folding is expected to provide further insights (Spitale and Incarnato 2022).

The set of canonical base pairs, including Watson-Crick-Franklin AU and GC base pairs and wobble GU base pairs, constitute the secondary structure for an RNA. For functional RNAs, the secondary structure tends to remain evolutionarily conserved for the function to be conserved. Improved RNA secondary structure prediction is key to discovering new functional RNAs (Gao et al. 2021; Fu et al. 2015; Gruber et al. 2010; Zhang et al. 2023) and offers a better understanding of the known ones (Hagey et al. 2022; Li et al. 2021; Szutkowska et al. 2022).

Comparative sequence analysis is the gold standard for RNA secondary structure determination in the absence of a solved 3D structure (Gutell et al. 2002). However, the availability of a large number of homologous sequences is a prerequisite for RNA secondary structure determination. Base pairs are identified by finding covariations in an alignment that is refined to reflect the structure. These covariations, for example an AU base pair in one sequence that replaces a GC base pair in another sequence, provide evidence for conserved pairs. Covariations occur frequently across evolution because RNA structure is conserved although sequence is not. The process of comparative analysis is semi-automated, but still requires considerable manual effort to deduce covariation and iteratively refine the predicted structure (Gutell 2014). One important reason for the difficulty in automating this process is that the covariations that reveal the conserved structure complicate the alignment of the sequences based on nucleotide identity alone.

The nearest neighbor thermodynamic model (a.k.a. Turner Rules) can be used to estimate the Gibbs free energy change of folding for a given secondary structure (ΔG°)(Xia et al. 1998; Mathews et al. 2004; Mittal et al. 2024). Dynamic programming algorithms using this model can predict the structure for a given sequence with reasonable accuracy by estimating the lowest free energy structure (Mathews et al. 2010). Dynamic programing algorithms are also used to estimate base pairing probabilities with partition function calculations (McCaskill 1990). The pairing probabilities offer a more nuanced view of the secondary structure, especially for sequences that sample multiple secondary structures (Mathews 2006). The base pairing probabilities also provide information about the confidence of predicted pairs (Mathews 2004).

Algorithms have been developed to automate comparative sequence analysis by applying structure prediction to a set of homologous sequences to predict the consensus structure that is the conserved secondary structure common to all sequences. ‘Align-then-fold’ has been one of the popular approaches to determine the consensus secondary structure using a set of homologous sequences (Bernhart et al. 2008; Hofacker et al. 2002; Knudsen and Hein 1999, 2003; Rivas 2020; Seeman et al. 2008). In this approach, a fixed input alignment is provided by the user. The thermodynamic nearest neighbor model along with covariation analysis is used to predict structures where base pairs are common for all sequences in the alignment. Dynamic programming is used to search for the best scoring consensus structure or the consensus base pairing probabilities. These approaches are popular because they work well when the sequences are similar and the computational cost is lower than approaches that simultaneously predict structure and the sequence alignment. The structure prediction accuracy is improved over single sequence structure prediction because structures with conserved base pairs are scored more favorably by the nearest neighbor model and the covariation terms.

As summarized in Figure 1, we report two new programs: AlignmentFold and AlignmentPartition. These take the align-then-fold approach to determine the consensus minimum free energy structure and base pairing probabilities, respectively. For this work, we developed principled free energy change parameters that account for gaps and non-canonical base pairing that arise during simultaneous folding of sequences (Mittal et al. 2024). We take advantage of the ability of RNAstructure to use an extended nucleotide alphabet, including the gap symbol, to use these new parameters (Kierzek et al. 2022).

**Figure 1.**
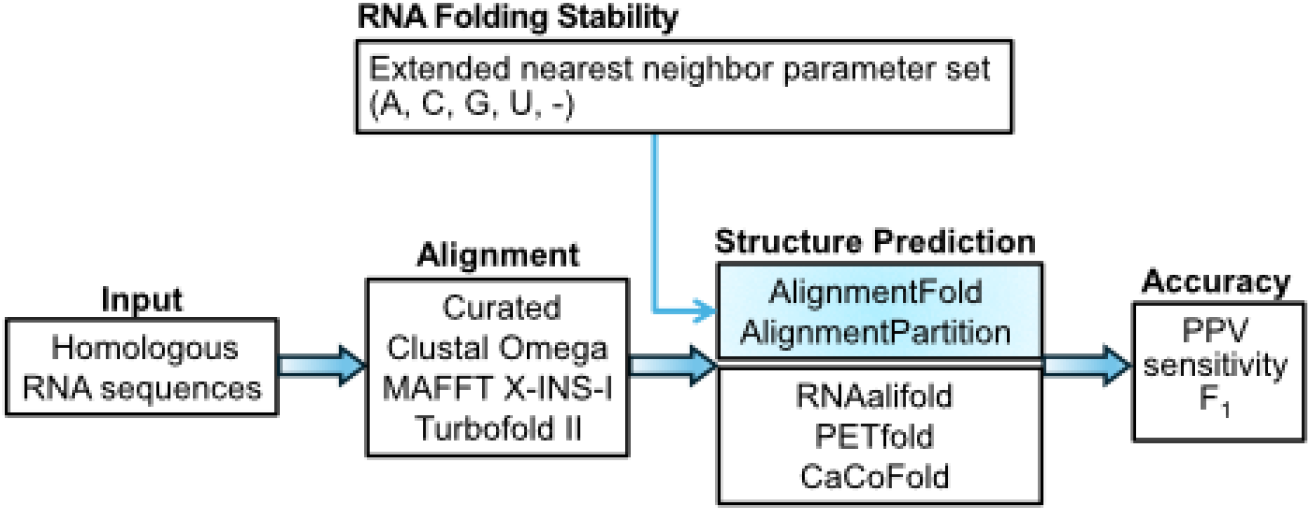
AlignmentFold and AlignmentPartition use an extended nearest neighbor parameter set to predict a consensus secondary structure from a fixed alignment of homologous RNAs. We derived principled parameters to include the gap symbol. In this work, we benchmarked structure prediction methods using manually curated alignments and predicted alignments. The predicted alignments test expected performance of these programs before the structure of an RNA family is known; the manually curated alignments are informed by structure.

We benchmarked our new programs against popular available programs, RNAalifold (Hofacker et al. 2002; Bernhart et al. 2008), PETfold (Seeman et al. 2008) and CaCoFold (Rivas 2020) using 5, 10, and 20 sequence sets designed to mimic the early stages of structure determination for newly discovered ncRNAs. In contrast to our expectations, we found that including terms that account for covariation analysis brings only modest improvement at best in prediction accuracy. An increase in the number of sequences in the input might bring marginal improvement in accuracy. We also tested the accuracy when using alignments generated by several sequence alignment programs. The alignments generated by MAFFT X-INS-I (Katoh and Toh 2008), which considers structure when aligning the sequences, lead to more accurate consensus structure prediction.

## RESULTS AND DISCUSSION

### Folding Model

AlignmentFold and AlignmentPartition find the lowest free energy secondary structure and partition function, respectively, by mapping predicted base pairs to all the sequences in a sequence alignment. The partition function estimates the pairing probabilities for base pairs in the conserved structure. The programs calculate the mean folding free energy change across the sequences for a given structure. The alignment gap symbol is treated as a nucleotide in an extended alphabet of bases. Base pairs other than AU, GC and, GU occur when the structure is mapped to the individual sequences in the alignment. These include non-canonical pairs like “GG” and gapped pairs like “-A” (Figure 2A). We developed nearest neighbor parameters to handle these cases, following the 2004 Turner Rules (Mathews et al. 2004; Mittal et al. 2024). This takes advantage of the ability to use extended alphabets of nucleotides (Kierzek et al. 2022). We also modified the single sequence folding software in the RNAstructure package to work with multiple sequence alignments. This provides suboptimal structure prediction (alternative low free energy structure prediction), simplifies maintenance and simplifies the incorporation of new features. The details of the implementation are provided in the METHODS section.

**Figure 2.**
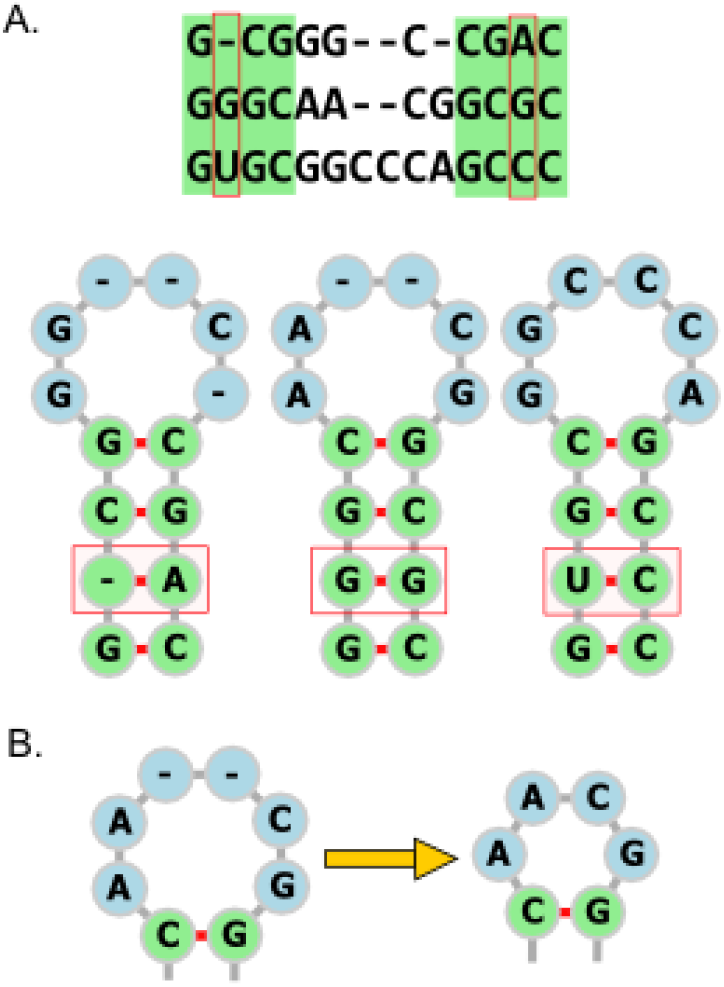
Appearances of alignment gaps and non-canonical pairs. Panel A shows a hairpin-stem-loop structure mapped to individual sequences in an alignment. This mapping can result in gaps appearing in base paired positions, for example the -A base pair that appears in the first sequence. The mappping results in a GG base pair in the second sequence. Panel B illustrates that gaps are removed from hairpin loops (and also internal loops) before free energy estimation. In this example, the six-membered hairpin loop contains two gaps that are removed to make a tetraloop.

### Parameter Optimization

To optimize four adjustable parameters (introduced below), we adopted a set of training RNA families that are separate from families used to test performance (Tan et al. 2017; Szikszai et al. 2022; Lu et al. 2009; Liu et al. 2010; Rivas et al. 2012). Consistent with our prior work, we used 5S ribosomal RNA (Eubacteria subfamily), group I intron (IC1 subfamily), tmRNA, and tRNA families for training and 16S rRNA (Alphaproteobacteria subfamily), SRP RNA (Protozoan subfamily), RNase P RNA (bacterial type A subfamily) and telomerase RNA families for testing, all of which were drawn from our RNAStralign data set (Tan et al. 2017).

For determining a consensus secondary structure from a fixed alignment, a criterion to determine positions that can base pair is required. For the alignment, positions are columns in the alignment, represented by nucleotides or gaps in each sequence. For AlignmentFold, we used a simple base pairing criterion where positions (i,j) can base pair if the fraction of sequences that can form a canonical base pair exceeds a cutoff. A higher, more restrictive cutoff leads to a higher positive predictive value (PPV; fraction of predicted pairs that are correct), lower sensitivity (fraction of known pairs correctly predicted), lower F_1_ score (the harmonic mean of sensitivity and PPV), and substantially improved run-time performance (Supplementary Table S1). To strike a balance between sensitivity, specificity, and runtime, we selected 0.3 as the default cutoff. A less restrictive base pairing cutoff can be chosen by the user to allow a greater number of pairs of positions in alignment to be eligible for base pairing. This is more resilient to alignment errors, but substantially increases the run time due to the expanded consensus secondary structure space.

The maximum size of an internal loop is set to 30 unpaired bases by default for single sequence folding in the RNAstructure package. For AlignmentFold, this was increased to 40 to accommodate the plausible increase in the size of internal loops due to the presence of gaps in the alignment. Increasing the maximum size of internal loops leads to improved sensitivity (Supplementary Table S2). It also leads to higher runtime due to the expanded consensus secondary structure space.

For AlignmentPartition, we used the same parameters, i.e. 0.3 as the base pairing cutoff and 40 as the maximum size of internal loop. This is important because the estimated pair probabilities reflect the same landscape of possible structures that free energy minimization structure prediction considers.

The ProbKnot (Bellaousov and Mathews 2010; Zhang et al. 2019) and MaxExpect (Lu et al. 2009) tools in the RNAstructure package predict structures from base pairing probabilities. We adapted these to use the output of AlignmentPartition to predict conserved structures. ProbKnot is capable of predicting pseudoknots (Liu et al. 2010), although AlignmentFold and MaxExpect are not. We found 4.0 to be the optimal value of the gamma parameter for MaxExpect on the training set (Supplementary Table S3) and 0.1 to be optimal threshold for ProbKnot (Supplementary Table S4).

### Accuracy Benchmarks

We benchmarked AlignmentFold, AlignmentPartition (generating structures with MaxExpect or ProbKnot), RNAalifold (Hofacker et al. 2002; Bernhart et al. 2008), PETfold (Seeman et al. 2008) and CaCoFold (Rivas 2020) on alignments estimated using Clustal Omega (Sievers et al. 2011), MAFFT X-INS-I (Katoh and Toh 2008) and TurboFold II (Tan et al. 2017) and also on the manually curated alignments. It is important to benchmark align-then-fold structure prediction tools using predicted alignments because the accepted alignments were generated using the structure for guidance (Malik et al. 2024). RNA secondary structure prediction from an alignment of homologous RNAs presents a chicken-and-egg problem where we need a high-quality sequence alignment to arrive at accurate prediction of conserved structure and _*vice-versa*_ (Seetin and Mathews 2012). Also, we used alignments of 5, 10, and 20 sequences to represent the early stages of structure determination when a limited number of homologous RNAs are available for a newly discovered RNA family.

The best performance for structure prediction accuracy was obtained using curated alignments, as expected because those are informed by the conserved structure (Supplementary Table S5). The best structure prediction accuracies for predicted alignments were obtained using MAFFT X-INS-i (Figure 3 and Supplementary Table S5). Prediction accuracy on sequence alignments generated using TurboFold II was similar (within 3%; Supplementary Tables S5-S9), except for SRP RNA where MAFFT X-INS-i is substantially better (Supplementary Table S8). Structure prediction accuracy is worst with alignments generated by Clustal Omega, which estimates alignments without considering the structure and relies solely on sequence identity. Also, PETfold performs substantially better (> 10%) on MAFFT X-INS-i-aligned RNase P sequences. Both MAFFT X-INS-i and TurboFold II consider secondary structure information generated through thermodynamic nearest neighbor-based computational approaches.

**Figure 3.**
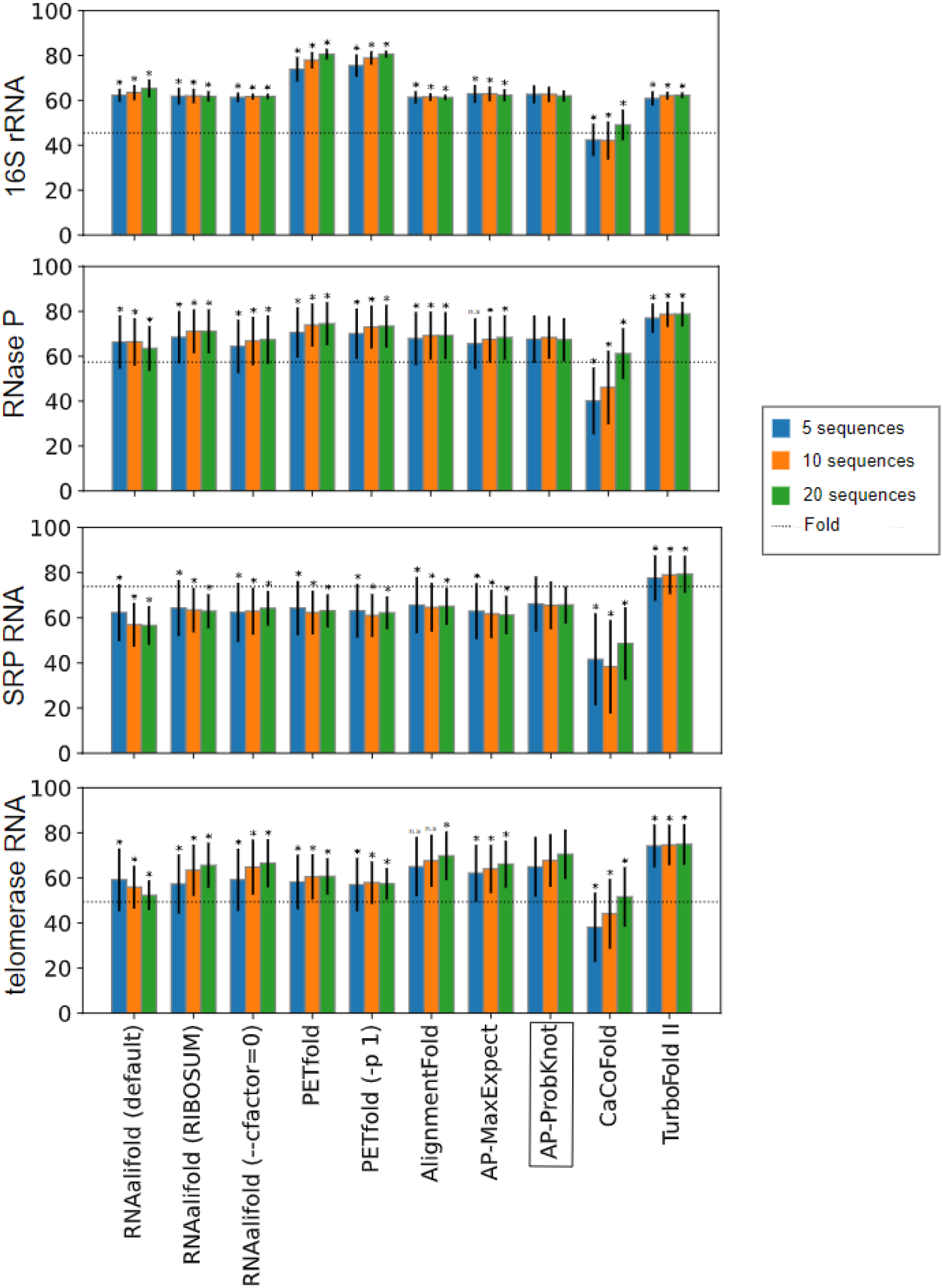
F_1_ score for RNAalifold (default), RNAalifold (RIBOSUM), RNAalifold (−-cfactor=0), PETfold, PETfold (−p 1), AlignmentFold, AlignmentProbability MaxExpect (AP-MaxExpect), AlignmentProbability ProbKnot (AP-ProbKnot), and TurboFold II on 200 sets of 5, 10 and 20 sequences each aligned using MAFFT X-INS-I except TurboFold II, which takes unaligned sequences as input. Plotted uncertainty is ± the standard deviation. The average F_1_ score of single sequence folding with RNAstructure Fold (lowest free energy structure prediction) has been plotted using a dotted line. F_1_ scores which are significantly different (p < 0.05) than AP-ProbKnot are marked by *.

The accuracies of most align-then-fold tools are similar, although there are idiosyncrasies observed across families. AlignmentFold, AlignmentPartition (ProbKnot and MaxExpect), PETfold and RNAalifold offer better prediction accuracy as measured by F_1_ score than single sequence structure prediction by Fold for 16S rRNA, RNase P and telomerase RNA (Figure 3). PPV and sensitivity are available as Supplementary Figures S1 and S2. For SRP RNA, these tools can offer superior PPV but substantially poorer sensitivity with structurally informed alignments (Supplementary Table S10). Except for CaCoFold, improvement in prediction accuracy in terms of F_1_ score with increase in the number of sequences is not guaranteed (Supplementary Table S5). The AlignmentPartition-ProbKnot structure predictions outperform all other methods including PETfold, RNAalifold, and CaCoFold for SRP RNA and telomerase RNA across all alignment modalities (Supplementary Figures S1 and S2 and Supplementary Tables S8 and S9). AlignmentPartition-MaxExpect predictions are more accurate than AlignmentFold and AlignmentPartition-ProbKnot for 16S rRNA (Figure 3 and Supplementary Table S6).

‘Align-and-fold’ algorithms, unlike ‘align-then-fold’ algorithms, take unaligned homologous sequences as input and return structures specific to each sequence in input along with an alignment. These algorithms tend to have better prediction accuracy than align-then-fold algorithms (Tan et al. 2017). TurboFold II is one such ‘align-and-fold’ algorithm. TurboFold II has better prediction accuracy than PETfold, RNAalifold, and AlignmentFold when input to these algorithms is an estimated alignment (Supplementary Table S11). The only exception being the 16S rRNA family where PETfold outperforms TurboFold II, and other align-then-fold methods also offer comparable accuracy to TurboFold II (Supplementary Tables S6 and S11). This is consistent with prior benchmarks where it was noted that the pairwise sequence identities of the 16S rRNA sequences is higher than that of other families (Tan et al. 2017)(Supplementary Figure S3).

### Comparison of Methods

In this work, we benchmarked RNAalifold version 2.4. It by default uses covariation-based scoring as described in 2002 (Hofacker et al. 2002). It uses a generalized formula to calculate the contribution of the covariation to the final score. RNAalifold with the -r option uses specialized RIBOSUM-like matrices for calculating the contribution of covariation to the final score as described in 2008 (Bernhart et al. 2008). It adapts to the levels of sequence identity in the alignment in that it stabilizes the formation of base pairs with compensatory mutations more generously when the level of sequence identity is high. The contribution of the covariation score to the total score often exceeds 50% (Supplementary Table S12).

RNAalifold allows the user to tune the contribution of covariation-based scoring _*vis-à-vis*_ thermodynamics for consensus structure prediction. A user can set it with ‘--cfactor’ option. By default, it is set to 1. We set it to 0 to benchmark RNAalifold without covariation-based scoring. Typically, RNAalifold with RIBOSUM-like scoring (−r flag) performs the best for the RNAalifold methods. It is closely followed by RNAalifold without covariation-based scoring, which in turn performs better than RNAalifold with default settings (Figure 4 and Supplementary Table S5).

**Figure 4.**
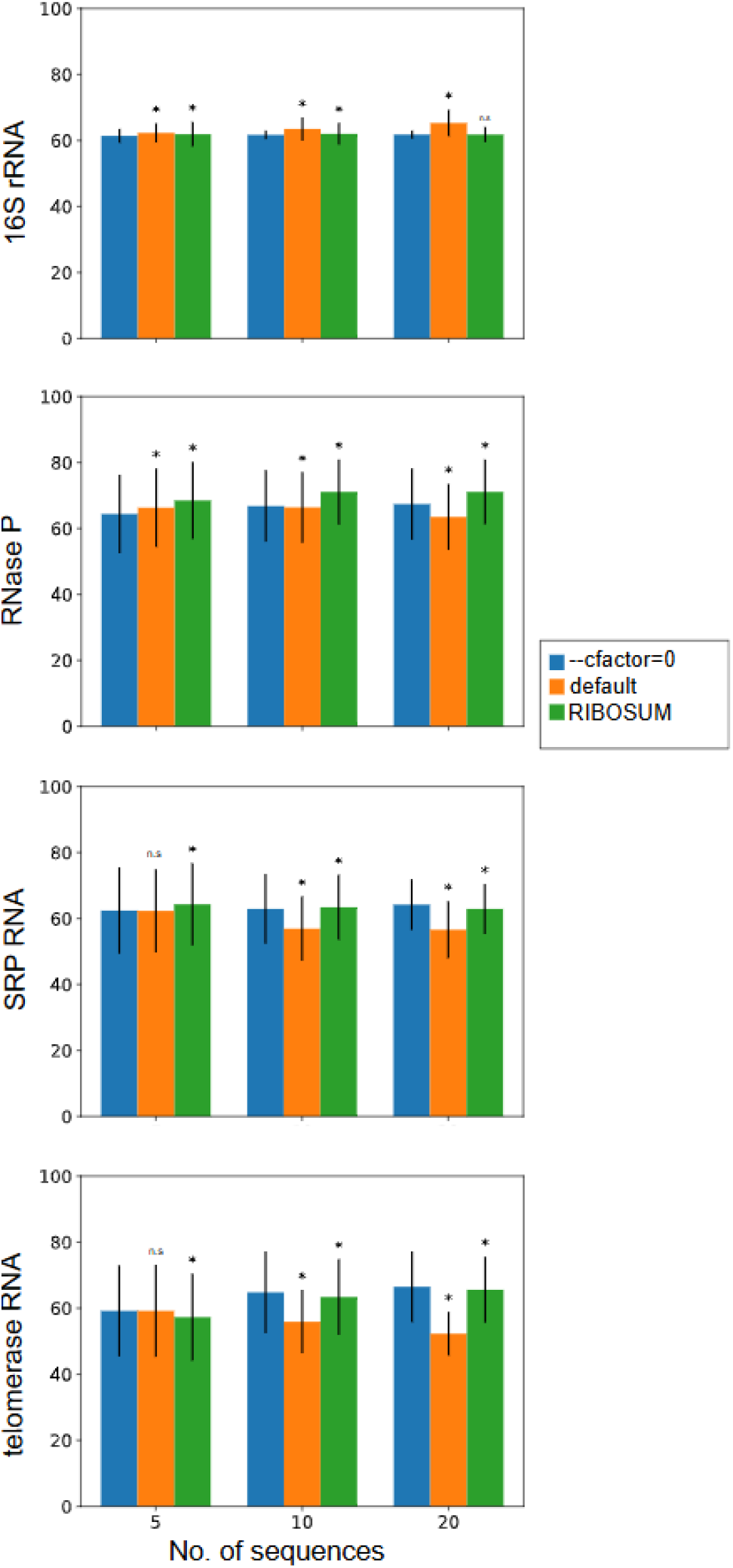
F_1_ score for RNAalifold (−-cfactor=0), RNAalifold (default), and RNAalifold (RIBOSUM) on 200 sets of 5, 10 and 20 sequences each aligned using MAFFT X-INS-i. Uncertainties are plotted as ± the standard deviation. F_1_ scores which are significantly different (p < 0.05) than RNAalifold (−-cfactor=0) are marked by *.

The F_1_ accuracy of RNAalifold (default) tends to decrease with increasing number of sequences in the input alignment (Supplementary Table S5). There are some exceptions to this trend. For example, RNAalifold (default) outperforms RNAalifold with no covariation-based scoring and RNAalifold with RIBOSUM-like scoring on the 16S rRNA family across all alignment methods (Supplementary Table S6). This is particularly interesting because RIBOSUM-like matrices were derived from analysis of 16S rRNA sequences (Bernhart et al. 2008).

PETfold provides a maximum expected accuracy structure by optimally combining pairing probabilities determined using RNAfold and Pfold (Knudsen and Hein 2003). While RNAfold relies on thermodynamic parameters to find the minimum free energy structure, Pfold draws from an explicit evolutionary model and SCFG-based probabilistic model for secondary structures. PETfold was optimized over curated alignments from Rfam.(Griffiths-Jones et al. 2005) The default parameter setting limits the influence of the evolutionary model on structure prediction and emphasizes the thermodynamics. PETfold removes columns with more than 25% gaps from the alignment before structure prediction and introduces a gap in their positions in the final consensus structure. It calculates “expected overlap” for a given consensus structure over all the sequences individually and avoids the issue of non-canonical pairing that arises during simultaneous folding of the sequences. PETfold with the -p option set at 1 relies solely on base pairing probabilities generated with RNAfold and does not consider probabilities generated by Pfold. Therefore, PETfold with -p 1 option does not consider covariation information. PETfold with default settings tends to perform slightly (∼1%) better in F_1_ score than PETfold with the -p 1 option (Supplementary Table S5; MAFFT X-INS-i alignments). However, there is an exception to this trend. PETfold without covariation information outperforms PETfold with default settings as well as all other methods on 16S rRNA alignments of size 5, 10, and 20 sequences across all alignment methods (Supplementary Table S6). But PETfold with default settings again performs better on large 16S rRNA alignments of 100 sequences (Figure 5 and Supplementary S13). PETfold with default settings amongst all methods offers the best F_1_ performance for 100 RNase P RNA sequences aligned with MAFFT X-INS-i or Clustal Omega (Supplementary Table S13 and Figure 5). PETfold offers suboptimal consensus structure prediction as well.

**Figure 5.**
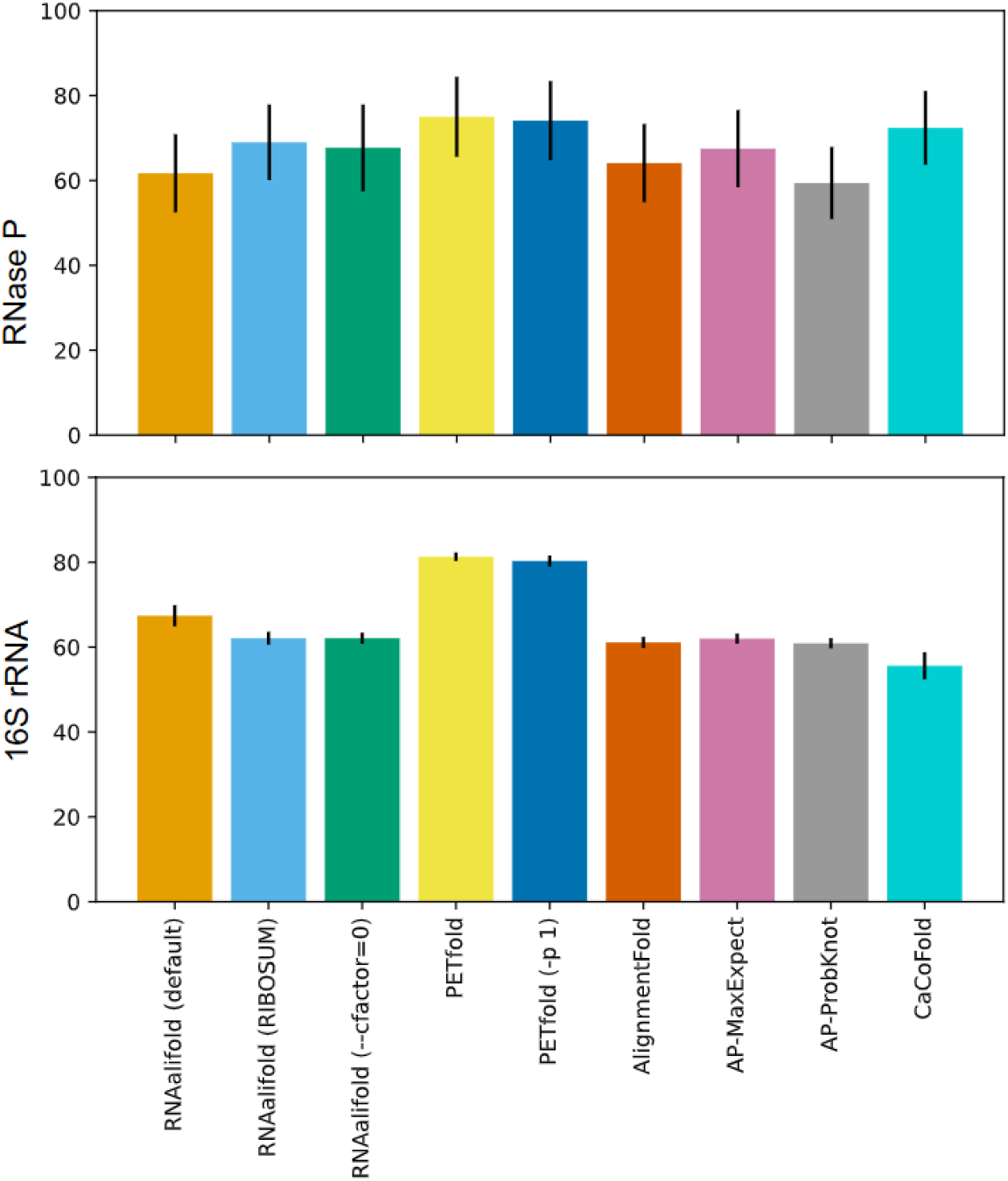
F_1_ score for RNAalifold (default), RNAalifold (RIBOSUM), RNAalifold (−-cfactor=0), PETfold, PETfold (−p 1), AlignmentFold, AlignmentProbaility MaxExpect, AlignmentProbability ProbKnot, and CaCoFold on 10 sets of 100 sequences each aligned using MAFFT X-INS-i.

CaCoFold (Cascade variation/covariation Constrained Folding algorithm) takes a three-fold approach to consensus structure prediction (Rivas 2020). First, it identifies pairs of positions in alignment for which covariation is statistically significant when considering phylogeny. Second, it identifies pairs of positions for which there is negative evidence of base pairing. These positions vary without significant covariation with other positions and are therefore forbidden from base pairing. Finally, folding is performed with probabilistic algorithms in a way that incorporates the above information about base pairing into one structure. Other methods benchmarked here consistently outperform CaCoFold on estimated alignments. CaCoFold, however, offers highest prediction accuracy for the curated alignment of RNase P RNA of 20 and 100 sequences (S7 and S13, Supplementary Information). These results suggest that CaCoFold requires a larger set of sequences to identify significant covariations. CaCoFold can predict pseudoknots like AlignmentPartition-ProbKnot and in contrast to RNAalifold, PETfold, AlignmentFold, and AlignmentPartition MaxExpect, which cannot predict pseudoknots.

### Time Benchmarks

The runtime complexity of AlignmentFold is O(HN^3^) where H is the number of sequences in the alignment and N is the length of alignment. This is comparable to O(N^3^) of single sequence folding where N is the length of the sequence. Supplementary Table S14 shows the runtime performance of AlignmentFold and AlignmentPartition.

## CONCLUSIONS

These benchmarks highlight the idiosyncratic performance landscape of align-then-fold methods across alignment prediction methods and sequence numbers. It also highlights the importance of benchmarking these tools on estimated alignments and across a range of number of sequences.

An important finding of these benchmarks on up to 20 sequences is that there is limited improvement offered by the inclusion of covariation information in structure prediction, especially for estimated alignments. They suggest that the accuracy is a result of a complex interplay of multiple factors like base pairing acceptance criteria, handling of gaps, and underlying folding models in secondary structure prediction from the fixed alignment. This work brings forth the advantages and limitations of align-then-fold tools vis-à-vis single sequence folding and align-and-fold approaches like TurboFold II.

AlignmentFold and AlignmentPartition offer an intuitive and transparent approach to consensus structure prediction. By using recursions previously developed for single sequence folding for processing multiple sequence alignments, it demonstrates an innovative approach towards software development for structure prediction. AlignmentFold and AlignmentPartition are incorporated in the RNAstructure software package, which is available freely under the GNU Public License at https://rna.urmc.rochester.edu.

## METHODS

### Software Implementation

The existing recursions in the RNAstructure package for single sequence folding were used for AlignmentFold and AlignmentPartition (Reuter and Mathews 2010). This was implemented with C++ function pointers. Function calls that calculate free energy increments for stacks and loops for a single sequence were replaced with function pointers. The pointers indicate functions that calculate these energies for multiple sequences when the input is a multiple sequence alignment and for a single sequence when the input is a single sequence. The energy model is therefore the full Turner 2004 model including the coaxial stacking of helices in multibranch and exterior loops (Mittal et al. 2024; Ward et al. 2019). This approach will facilitate updates to the software, for example, as the free energy model undergoes further improvements (Zuber et al. 2022).

### Improving run time performance

For a given alignment, a 2D matrix stores information about whether the positions i,j in the alignment are eligible for base pairing as per the criteria. Another set of 2D matrices store information about the number of gaps between any two positions for each sequence in the alignment. For each sequence in the alignment, a version of the sequence without gaps is also stored. A 1-D vector corresponding to each sequence in the alignment serves as a map for base positions from a sequence in alignment with gaps to the gapless sequence. All the above-mentioned entities are initialized at the beginning of execution, and these allow rapid look up during run time leading to faster execution.

### Benchmarking run time performance

For benchmarking run time performance, AlignmentFold and AlignmentPartition were compiled using GCC 4.8 and run on Red Hat Enterprise Linux 7 Maipo (x86_64) with an Intel Xeon CPU E5-2695 v2 @ 2.40GHz and 4 GB memory.

### Determining Structure Prediction Accuracy

The accuracy of RNA secondary structure prediction is calculated in terms of PPV (positive predictive value), sensitivity, and F_1_ score. PPV is the fraction of the predicted base pairs that also occur in the accepted structure. Sensitivity is the fraction of base pairs in the accepted structure that are correctly predicted. F_1_ score is harmonic mean of sensitivity and PPV (Mathews 2019).

Predicted base pairs are considered correct if a nucleotide either on 5’ or 3’ end of the correct base is off by one position (Mathews et al. 1999). For instance, a predicted base pair (i, j) is correct if base pair (i, j), (i±1, j) or (i, j±1) exists in the accepted structure. This is due to uncertainty in secondary structure determination by comparative sequence analysis (Gutell et al. 2002) and thermodynamic fluctuation in local structures (Fu et al. 2014; Znosko et al. 2002; Woodson and Crothers 1987). The scorer program of RNAstructure was used to calculate PPV and sensitivity.

### Benchmark Data

The RNAStralign (Tan et al. 2017) database was divided and used for determining hyperparameters and for accuracy benchmarking. RNAStralign is aggregated from expertly curated online databases of RNA secondary structure and sequence alignment. Sequences and their structures in RNAStralign are categorized into homologous families. The families included are 5S ribosomal RNA (Szymanski et al. 2002), Group I intron (Zhou et al. 2008), tmRNA (Zwieb et al. 2003), tRNA (Jühling et al. 2009), 16S ribosomal RNA (Cannone et al. 2002), Signal Recognition Particle (SRP) RNA (Rosenblad et al. 2003), RNase P RNA (Brown 1999), and telomerase RNA (Nawrocki et al. 2015). 16S rRNA, 5S rRNA, RNase P RNA, SRP RNA and group I intron families are further categorized into subfamilies in line with the original databases.

### Optimizing base pairing cutoff parameter

In AlignmentFold and AlignmentPartition, base pairs are only permitted when the fraction of sequences that can form canonical base pairs exceeds a pairing cutoff. To determine the optimal base pairing cutoff, 250 groups of input sequences, comprising five, ten and twenty homolog sets, were randomly selected from the 5S ribosomal RNA (Eubacteria subfamily), group I intron (IC1 subfamily), tmRNA, and tRNA families of the RNAStralign database. These sequence sets were aligned using MAFFT (X-INS-i)(Katoh and Toh 2008). A consensus structure was determined for each of the alignments using AlignmentFold with varied base pairing cutoff. Cutoffs 0.1, 0.2, 0.3, 0.4, 0.5, 0.6, 0.7 and 0.8 were benchmarked (Supplementary Table S1). The consensus structure was mapped back to each sequence in the alignment to determine the predicted structure for each sequence. Only AU, GC and GU base pairs were retained during mapping. Accuracy was calculated for each sequence individually. 0.3 was selected as the default base pairing cutoff for both AlignmentFold and AlignmentPartition.

### Optimizing maximum size of internal loop

In AlignmentFold and AlignmentPartition, there is a maximum number of unpaired nucleotides for internal and bulge loops, which is adjustable. Secondary structure space is not explored for internal loops of greater size for structure prediction. To determine the default size, alignments were generated as described for optimizing the base pairing cutoff above. A consensus structure was determined for each of the alignments using AlignmentFold with varied maximum internal loop sizes. Sizes 30, 35, 40, 45, 50, 55, 60 and 80 were benchmarked (Supplementary Table S2). The consensus structure was mapped back to each sequence in the alignment. Only AU, GC and GU base pairs were retained during mapping. Accuracy was calculated for each sequence individually. 40 was selected as the default maximum internal loop size for both AlignmentFold and AlignmentPartition.

### Optimizing gamma for MaxExpect

MaxExpect uses a partition function save file as input. This file contains the information needed for estimating base pairing probabilities (Lu et al. 2009). MaxExpect infers a structure by maximizing the sum of the base-pairing and single stranded nucleotide probabilities. Pairing probabilities are weighted by factor γ.

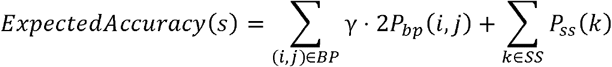

where *P*_*bp*_(*i,j*)is the base pairing probability for nucleotides at positions *i* and *j* and *P*_*ss*_(*k*) is the probability of remaining single-stranded (unpaired) for nucleotide at position *k*. The sums are over all base pairs (BPs) and single stranded nucleotides (SS) in structure *s*.

To determine the optimal gamma, alignments were generated from the training set as described above for optimizing the base pairing cutoff. Partition function save files were generated for alignments using AlignmentPartition. Maximum expected accuracy structures were generated for varied values of gamma: 0.5, 1, 2, 4, 5, 6, 7, and 8 (Supplementary Table S3). The structures were mapped back to each sequence in the alignment. Only AU, GC and GU base pairs were retained during mapping. Accuracy was calculated for each sequence individually. 4 was selected as the default γ for structure determination using MaxExpect.

### Optimizing threshold for ProbKnot

ProbKnot uses a partition function save file as input and returns a structure with the most probable base pairs. Each base pair (*i,j*) in this structure has probability *p*(*i,j*) that is highest amongst the competing pairs(Zhang et al. 2019). Also, *p*(*i,j*) is greater than or equal to a threshold probability.

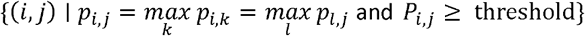

To determine the default threshold, alignments were generated from the training set as described earlier. Structures were generated with varied values of threshold: 0.0, 0.1, 0.2, 0.3, 0.4, 0.6, and 0.8 (Supplementary Table S4). The structures were mapped back to each sequence in the alignment. Only AU, GC and GU base pairs were retained during mapping. Accuracy was calculated for each sequence individually. 0.1 was selected as optimal threshold.

### Isolated base pairs

In single sequence folding, a base pair (i,j) is considered isolated if base pairs (i-1,j+1) and (i+1,j-1) cannot base pair. Single sequence folding in the RNAstructure package by default does not allow these isolated base pairs, but users have an option to allow them. In AlignmentFold and AlignmentPartition, base pairing between positions (i,j) is considered isolated if positions (i,j) fulfill the base pairing criterion stated above but neither (i+1,j-1) nor (i-1,j+1) positions do so. However, an exception is made and pairing is allowed for positions (i,j) when there exist positions (m,n) such that m can pair to n, i<m<n< j and all the intervening positions (i+1,i+2…m-2, m-1, n+1,n+2…j-2,j-1) have at least 50% nucleotides as gaps. Similarly, base pairing is allowed for (i,j) when there exist positions (m,n) such that m can pair to n, m<i and n>j and all the intervening positions (m+1,m+2…i-2, i-1, j+1,j+2…n-2,n-1) have at least 50% nucleotides as gaps.

### Accuracy Benchmarks

The command line options used for benchmarking each program are provided in Table 1.

**Table 1.** Commands for methods of sequence alignment and structure prediction benchmarked in this work.

| Method | Version | Command |
| --- | --- | --- |
| MAFFT with X-INS-i option | v7.490 (2021/Oct/30) | mafft-xinsi sequence_group.fasta |
| RNAalifold (default) | 2.4.14 | RNAalifold sequence_alignment.fasta |
| RNAalifold (no covariation) | 2.4.14 | RNAalifold --cfactor=0.0 sequence_alignment.fasta |
| RNAalifold with ribosum scoring | 2.4.14 | RNAalifold -r sequence_alignment.fasta |
| CaCoFold* | 2.0.4.a/ | rscape_v2.0.4.a/bin/R-scape --cacofold sequence_alignment.sto |
| AlignmentFold | 6.5 | RNAstructure/exe/AlignmentFold -b 0.3 -l 40 sequence_alignment.fasta consensus_structure.ct |
| AlignmentPartition | 6.5 | RNAstructure/exe/AlignmentPartition -b 0.3 -l 40 sequence_alignment.fasta pairing_probability.pfs |
| MaxExpect | 6.5 | RNAstructure/exe/MaxExpect pairing_probability.pfs mea_structure.ct -g 4 |
| ProbKnot | 6.5 | RNAstructure/exe/ProbKnot sequence_alignment.pfs mea_structure.ct -T 0.1 |
| TurboFold II | 6.5 | RNAstructure/exe/TurboFold config |
| Clustal Omega | 1.2.4 | clustalo -i sequence_group.fasta |
| PETfold | v2.2 | PETfold -f sequence_alignment.fasta |
| PETfold (no evolution) | v2.2 | PETfold -f sequence_alignment.fasta -p 1 |

\*CaCoFold takes input in Stockholm format

For benchmarking, 200 groups comprising of five, ten and twenty homologous sequences were randomly selected with replacement from the 16S rRNA (Alphaproteobacteria subfamily), SRP RNA (Protozoan subfamily), RNase P RNA (bacterial type A subfamily) and telomerase RNA. For SRP RNA, sequences shorter than 200 nucleotides were excluded because their structures are not consistent with those of longer sequences (Tan et al. 2017; Rosenblad et al. 2003). Similarly, for 16S rRNA sequences shorter than 1400 nucleotides were excluded. The benchmarks were therefore performed on families other than those used to train the pairing cutoff parameter (Szikszai et al. 2022; Rivas et al. 2012; Lu et al. 2009).

For benchmarks, the sequences were aligned using a multiple sequence alignment software or extracted from curated alignment and then consensus structure prediction was performed. The consensus structure was mapped back to each of the individual sequences and the accuracy determined. All methods were benchmarked on the same groups of sequences. As a control, single sequence folding was performed on each individual sequence using Fold (single sequence free energy minimization) and the accuracy calculated. Although a sequence can appear multiple times in sequence alignments, the accuracy of a sequence is averaged only once for the Fold calculations.

Benchmarking was also performed on 10 groups of 100 homologous sequences randomly selected with replacement from 16S rRNA (Alphaproteobacteria subfamily) and RNase P RNA (bacterial type A subfamily). The benchmarking was limited to sequences aligned using MAFFT X-INS-i and those extracted from curated alignments.

### Significance testing

To assess the statistical significance of the differences in sensitivity, PPV and F_1_, two-tailed, paired t-tests were performed using the ‘scipy.stats.ttest_rel’ function of SciPy v1.2.3 (Virtanen et al. 2020). The type I error rate was set to 0.05.

### Thermodynamic parameters for gaps and base pairs other than AU, GC, and GU

AlignmentFold maps the structure to all the sequences in the alignment while searching for the optimal structure. It treats gaps as a nucleotide. Base pairs other than AU, GC and, GU occur when the structure is mapped to the individual sequences in the alignment. We developed new nearest neighbor parameters to handle cases where the non-canonical pairs are predicted as base pairs or when gap symbols appear in pairs. In general, we average Turner 2004 parameters to estimate a free energy change for the resulting structure with the gap removed. This is intended to balance the accuracy of folding free energy estimates with computational efficiency.

We used the extended alphabet feature of RNAstructure to account for gaps, “-” as an additional nucleotide (Kierzek et al. 2022). In the RNAstructure package, all the files containing the energy parameters reside in the ‘data_tables’ directory. The files containing the extended parameter set start with ‘msa’.

Gaps are removed from hairpin loops and internal loops before free energy evaluation in AlignmentFold (See Figure 2B), following the strategy used in RNAalifold (Bernhart et al. 2008). If the unpaired number of nucleotides in the hairpin loop are fewer than 3, which is forbidden in single sequence folding, we use 8 kcal/mol as the loop closure penalty. This approximates the 5.4 kcal/mol cost of closing a hairpin of 3 unpaired nucleotides, plus the average cost of opening a base pair stack (2.1 kcal/mol). For a bulge or an internal loop, if the number of unpaired nucleotides is zero after removal of gaps in the loop region, we treat the motif as a stack of base pairs.

Helix stack parameters are specified in msa.stack.dg. When a gap-gap pair is followed by another gap-gap pair, a non-canonical pair, or a canonical pair, the stack is 0 kcal/mol. When a gap-nucleotide base pair is followed by a non-canonical pair, a canonical pair, or a gap-gap pair, 0.6 kcal/mol, the penalty used per nucleotide asymmetry in internal loops is used. When a gap-nucleotide base pair is followed by another gap-nucleotide base pair where the gaps appear in opposite strands, 0 kcal/mol is used. When a gap-nucleotide base pair if followed by another gap-nucleotide base pair where the gaps appear on the same strand, 1.2 kcal/mol is used because this is twice the asymmetry penalty.

When a non-canonical base pair follows another non-canonical base pair, 0.2 kcal/mol is used. This is equivalent to free energy change incurred by adding two unpaired nucleotides to an internal loop. For stacks consisting of one non-canonical base pair and one canonical base pair, the free energy change was estimated by linear regression from RNA tables for 1×1 internal loops and terminal mismatches. We treated the 1×1 internal loop table (rna.int11.dg) as combinations of two stacks consisting of one non-canonical base pair and one canonical base pair. Similarly, we treated the terminal mismatches (rna.tstack.dg) as a stack of a pair and a non-canonical pair. With these data, linear regression estimated the sequence dependent free energy contribution of the stack of a canonical pair followed by a non-canonical pair. These stacking parameters are also used for the coaxial stacking of adjacent helices with no intervening mismatch as contained in msa.coaxial.dg (Walter and Turner 1994).

For dangling ends (msa.dangle.dg), when the base pair adjacent to the dangling base is non-canonical or contains one or more gaps, 0 kcal/mol is used. Also, when the dangling base is a gap, 0 kcal/mol is used.

Terminal mismatch stacks are specified in msa.tstack.dg for exterior loops, in msa.tstackm.dg for multibranch loops, and in msa.tstackcoax for coaxial stacks of adjacent helices with an intervening mismatch (Tyagi and Mathews 2007; Kim et al. 1996; Walter et al. 1994). Following the 2004 nearest neighbor parameters, these sequence-dependent terminal mismatches values are identical in these different structural contexts (Mittal et al. 2024). When the terminal base pair is non-canonical pair or contains one or two gaps, 0 kcal/mol is used. If the terminal base pair is canonical and one of the mismatching bases is a gap, then the motif is treated as a dangling end and corresponding free energy from rna.dangle.dg is used. When both nucleotides in the terminal mismatch are gaps, 0 kcal/mol is used.

Internal loops of size 1×n (n>2), 2×3, or i×j (i,j >2) can be stabilized by first mismatches at each end (msa.tstacki.dg, msa.tstack1n.dg, and msa.tstack23.dg). When non-canonical base pairing or gaps are present in first mismatches, 0 kcal/mol is used. When one or both the mismatching bases are gaps, 0 kcal/mol is used as well.

First mismatches in hairpin loops are specified in msa.tstackh.dg. When the terminal base pair is non-canonical or contains gaps, 0 kcal/mol is used. If one of the mismatched bases is a gap, then the motif is treated as a dangling end and corresponding free energy from rna.dangle.dg is used. If both nucleotides in the mismatch are gaps, then 0 kcal/mol is used.

For the sequence-independent coaxial stack of two adjacent helices with an intervening mismatch (msa.coaxstack.dg), 0 kcal/mol is used when non-canonical base pair is followed by a mismatch or one or more bases in the terminal base pair or the mismatching bases are gaps. In these cases, the two helices would be beyond the single mismatch allowed for coaxial stacking.

Internal loops of size 2×2 are specified in msa.int22.dg. When one or more gaps appear in closing base pairs of 2×2 internal loops, then they are treated as “wildcards” (i.e. they are treated as though they could represent any of the canonical nucleotides A, C, G, or U) and the mean free energy change across all sequence possibilities for the gap is used.

When one or more non-canonical base pairs close a 2×2 internal loop, we treat this as a larger internal loop. We assume a canonical base pair adjacent to the non-canonical pair. For a single non-canonical pair closure, the free energy is the mean of canonical closures at that position with additional penalty of 0.2 kcal/mol, which would be incurred by a 2×2 internal loop becoming a 3×3 internal loop. When both the closing base pairs for the 2×2 internal loop are non-canonical, the free energy change is the mean of canonical closures with twice the 0.2 kcal/mol penalty. When gaps and non-canonical base pairing appear in closing base pairs for 2×2 loops, we treat gaps as “wildcards” and assume presence of a canonical base pair adjacent to the non-canonical pair. We use the mean free energy increment of all relevant possible sequences and canonical closures, with additional penalty of 0.2 kcal/mol. For all these cases, if the free energy change calculated is less than 0 kcal/mol, we use a minimum free energy change of 0 kcal/mol instead.

Internal loops of size 2×1 are specified in msa.int21.dg. When one or more gaps appear in closing base pairs of 2×1 internal loops, they are treated as “wildcards” and the mean free energy change across all sequence possibilities for the “wildcard” is used. When a non-canonical base pair closes 2×1 internal loop, we treat this as a larger internal loop. We assume a canonical base pair adjacent to the non-canonical pair. The free energy change is the mean of canonical closures at that position, with an additional penalty of 0.2 kcal/mol, which would be incurred by 2×1 loop becoming 3×2 loop. When both the closing base pairs for 2×1 internal loop are non-canonical, we make similar assumptions. The free energy change is the mean of canonical closures with twice the penalty. When gaps and non-canonical base pairing appear in closing base pairs for 2×1 loops, we treat gaps as “wildcards” and assume presence of a canonical base pair adjacent to the non-canonical pair. We take the mean of relevant possibilities and canonical closures, with an additional penalty of 0.2 kcal/mol. For all these cases, if free energy change calculated is less than 0 kcal/mol, we use 0 kcal/mol instead.

Internal loops of size 1×1 are parameterized in msa.int11.dg. When one or more gaps appear in closing base pairs of 1×1 internal loops, hey are treated as “wildcards” and the mean free energy across all sequence possibilities is used. When a non-canonical base pair closes a 1×1 internal loop, we treat this as larger internal loop. We assume a canonical base pair adjacent to the non-canonical pair. The free energy is the mean of canonical closures at that position, with additional penalty of 0.2 kcal/mol (which would be incurred by 1×1 loop becoming 2×2 loop). When both the closing base pairs for 1×1 internal loop are non-canonical, we make similar assumptions. The free energy change is the mean of canonical closures with twice the penalty. When gaps and non-canonical base pairing appear in closing base pairs for 1×1 loops, we treat gaps as “wildcards” and assume the presence of a canonical base pair adjacent to the non-canonical pair. We take mean of relevant possibilities and canonical closures, with an additional penalty of 0.2 kcal/mol. For all these cases, if free energy change calculated is less than 0 kcal/mol, we use 0 kcal/mol instead.

The specific hairpin loop sequence tables, including the tetraloop table (msa.tloop.dg), the triloop table (msa.triloop.dg), and the hexaloop table (msa.hexaloop.dg) remain identical to their tables in the 2004 nearest neighbor parameters. msa.miscloop.dg remains the same as rna.miscloop.dg. This includes the multibranch loop parameters and the helix end penalties. The length-dependent free energy penalties for closing hairpin, internal, and bulge loops (contained in msa.loop.dg) are sequence independent and remain the same as those in 2004 nearest neighbor parameters.

## ACKNOWLEDGEMENTS

This work was supported by United States National Institutes of Health grant NIH R35 GM145283 to D.H.M. We thank Liang Huang and Apoorv Malik (Oregon State University) for discussions.

