## Supplemental Tables and Figures for "AlignmentFold and AlignmentPartition: Improving the align-then-fold approach for RNA secondary structure prediction"

Table S1: AlignmentFold prediction accuracy (in percent) and run time (in seconds) as a function of base pairing cutoff. Sequences were aligned using MAFFT X-INS-i and maximum internal loop size was set to 40. For each family, 250 calculations were performed, choosing sequences at random. The mean across the 250 predictions is reported.

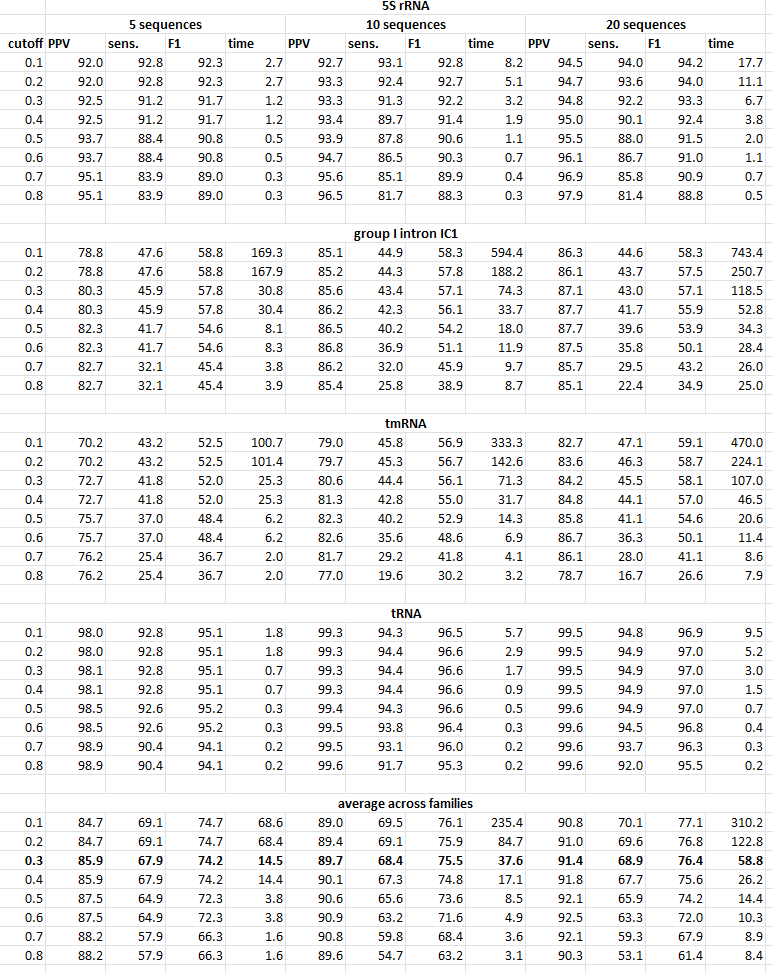

Table S2: AlignmentFold prediction accuracy (in percent) and run time (in seconds) as a function of maximum internal loop size. Sequences were aligned using MAFFT X-INS-i and base pairing cutoff was set to 0.3. For each family, 250 calculations were performed, choosing sequences at random. The mean across the 250 predictions is reported.

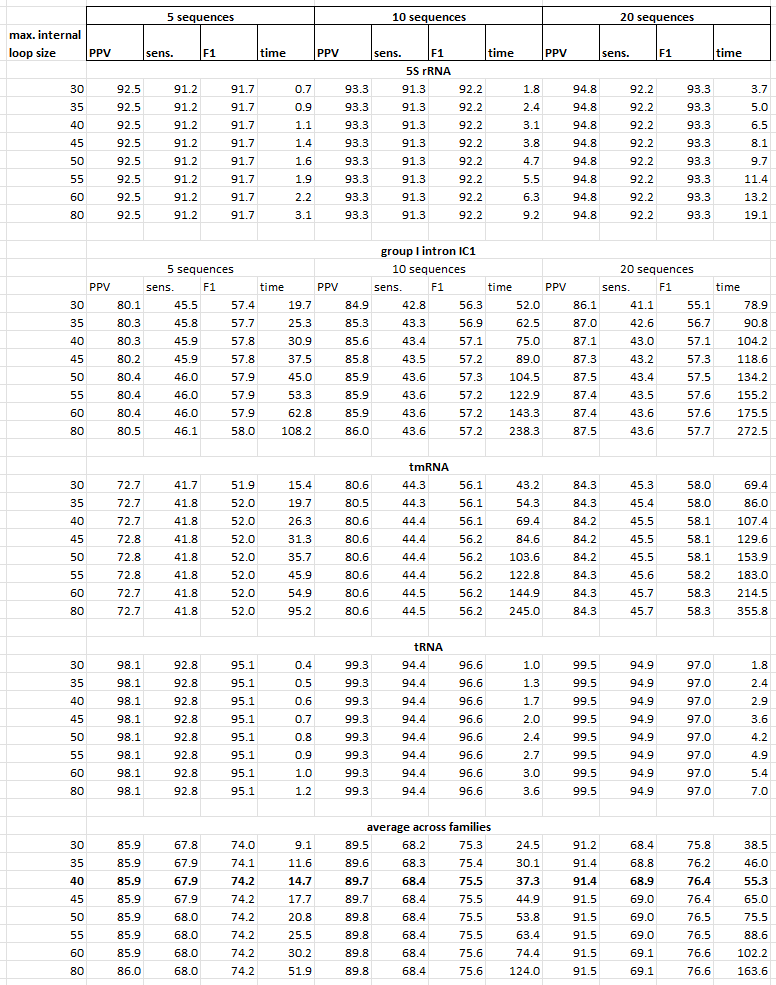

Table S3: Accuracy of MEA secondary structure prediction (in percent) as a function of gamma. Consensus base pairing probabilities were generated using AlignmentPartition for MAFFT X-INS-i alignments. The base pairing cutoff was set to 0.3 and maximum internal loop size was set to 40. MEA structure was determined from the consensus base pairing probabilities for various gamma values (0.5,1,2,4,5,6,7,8).

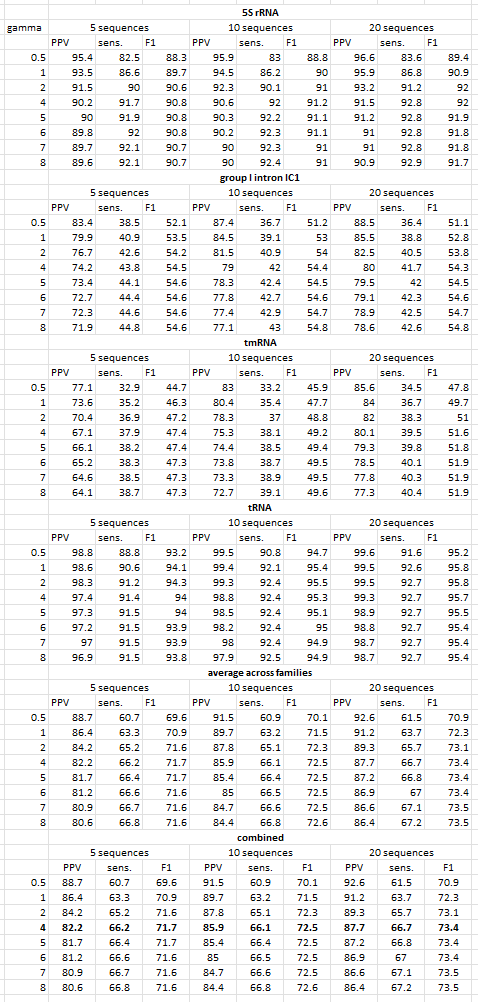

*Table S4: Prediction accuracy (in percent) for ProbKnot structures as a function of threshold. Consensus base pairing probabilities were generated using AlignmentPartition for MAFFT X-INS-i alignments. Base pairing cutoff was set to 0.3 and maximum internal loop size was set to 40. ProbKnot structures was determined from the consensus base pairing probabilities for various threshold (0.0, 0.1, 0.2, 0.3, 0.4, 0.6, 0.8).*

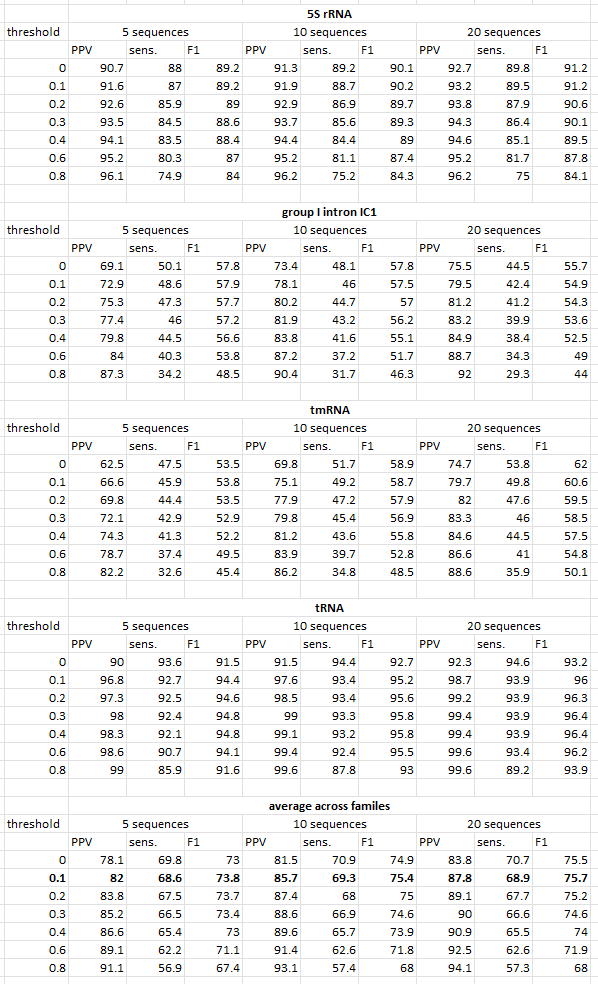

*Table* *S5: Average prediction accuracy (in percent) of align-the-fold methods across the four RNA families of the testing set (Tables S6-S9). Tables are provided for each alignment method, as indicated with the bold alignment method in the upper left corner. PPV, sensitivity, and F1 scores are provided for 5, 10, and 20 sequence calculations.*

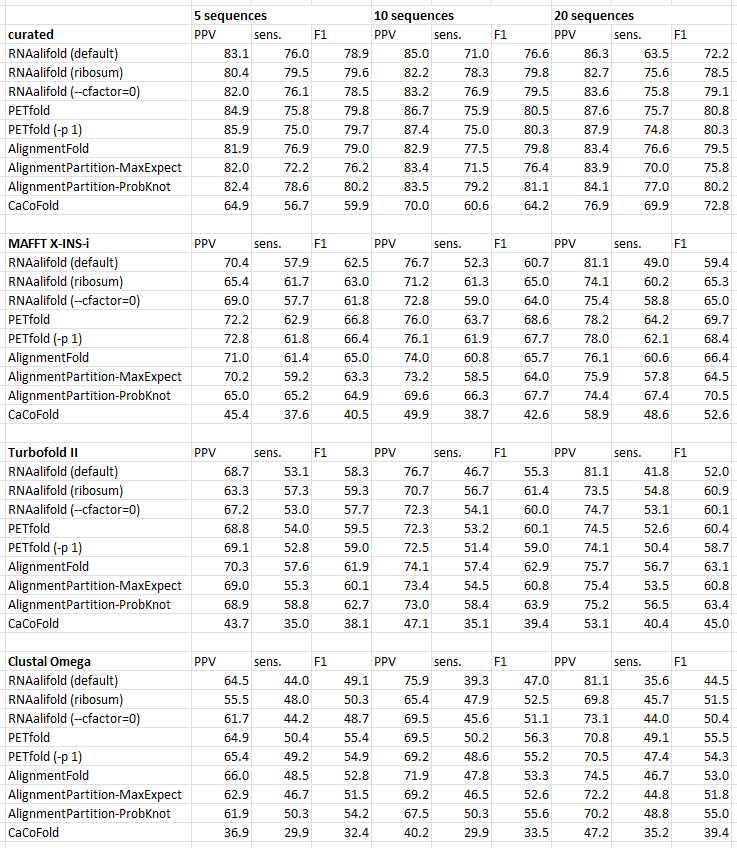

*Table* *S6: Prediction accuracy of align-the-fold methods for 16S rRNA. Tables are provided for each alignment method, as indicated with the bold alignment method in the upper left corner. PPV, sensitivity, and F1 scores are provided for 5, 10, and 20 sequence calculations.*

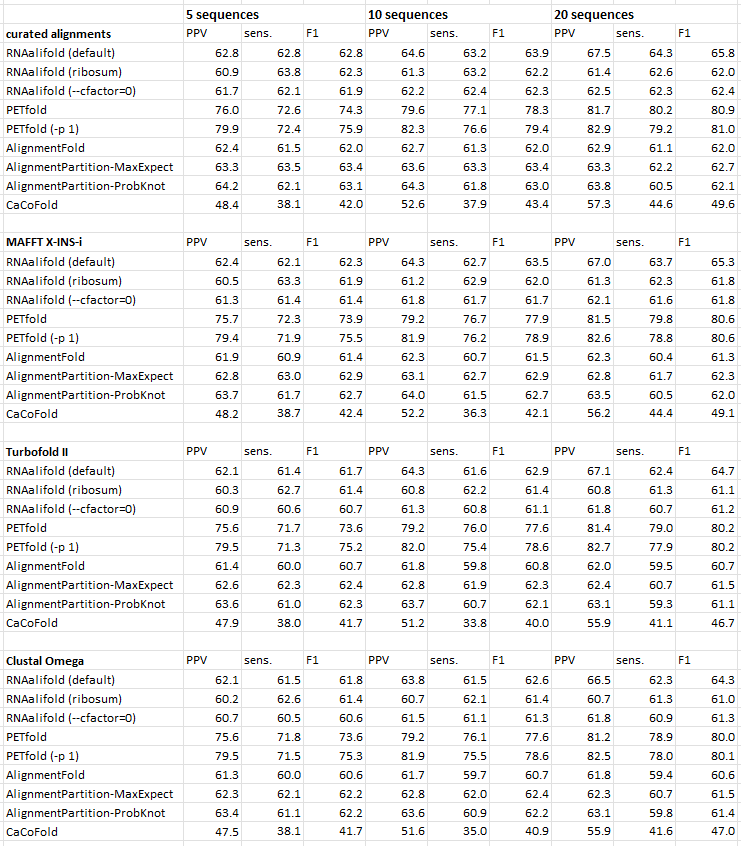

*Table* *S7: Prediction accuracy of align-the-fold methods for RNase P. Tables are provided for each alignment method, as indicated with the bold alignment method in the upper left corner. PPV, sensitivity, and F1 scores are provided for 5, 10, and 20 sequence calculations.*

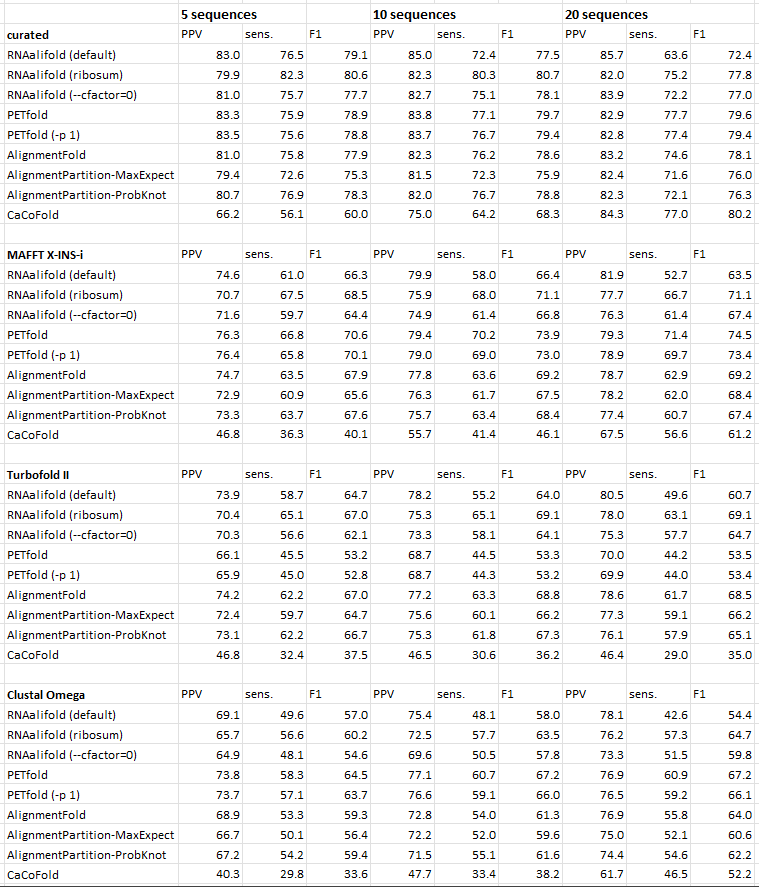

*Table* *S8: Prediction accuracy of align-the-fold methods for SRP RNA. Tables are provided for each alignment method, as indicated with the bold alignment method in the upper left corner. PPV, sensitivity, and F1 scores are provided for 5, 10, and 20 sequence calculations.*

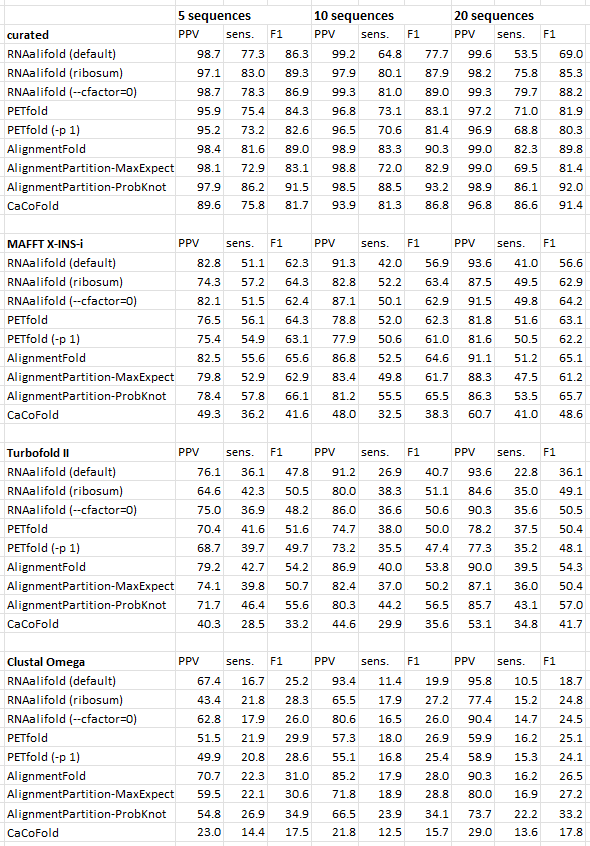

*Table* *S9: Prediction accuracy of align-the-fold methods for telomerase RNA. Tables are provided for each alignment method, as indicated with the bold alignment method in the upper left corner. PPV, sensitivity, and F1 scores are provided for 5, 10, and 20 sequence calculations.*

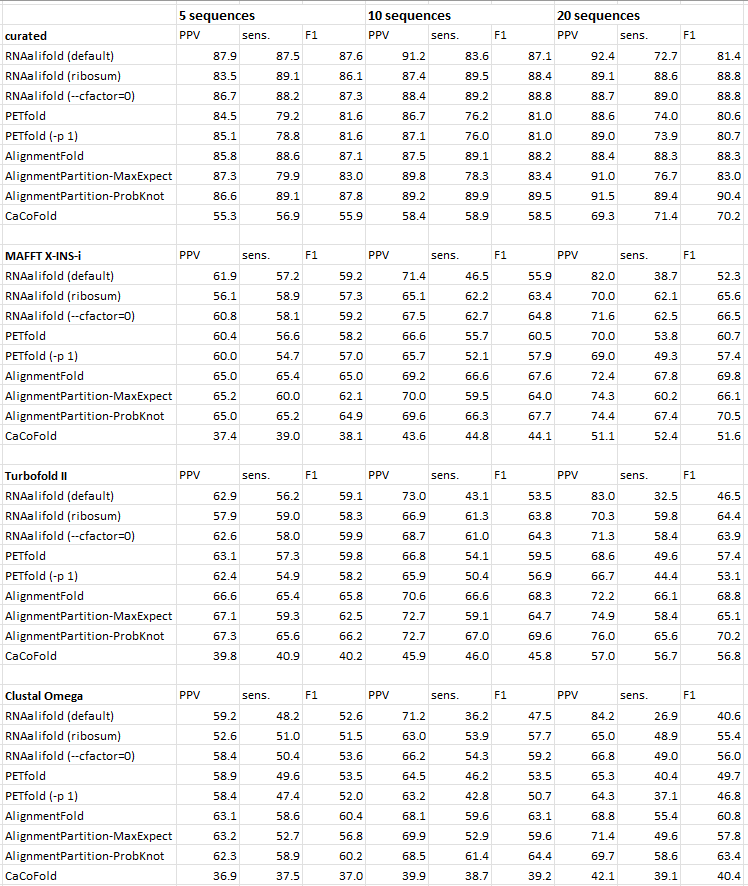

Table S10: Prediction accuracy of single sequence folding with Fold and partition.

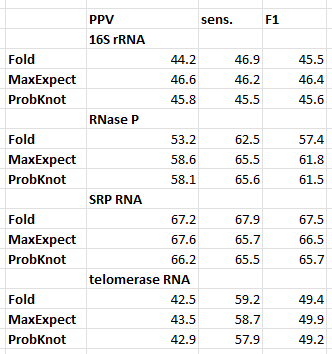

Table S11: Turbofold II prediction accuracy.

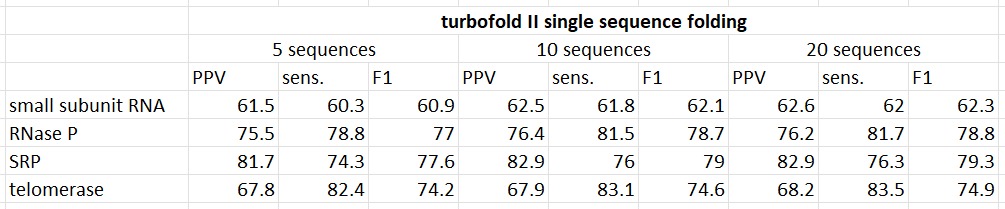

Table S12: Contribution of covariation towards final score in percent. RNAalifold reports contribution of covariation to the final score in kcal/mol. Average percent contribution of covariation to the final score is reported below.

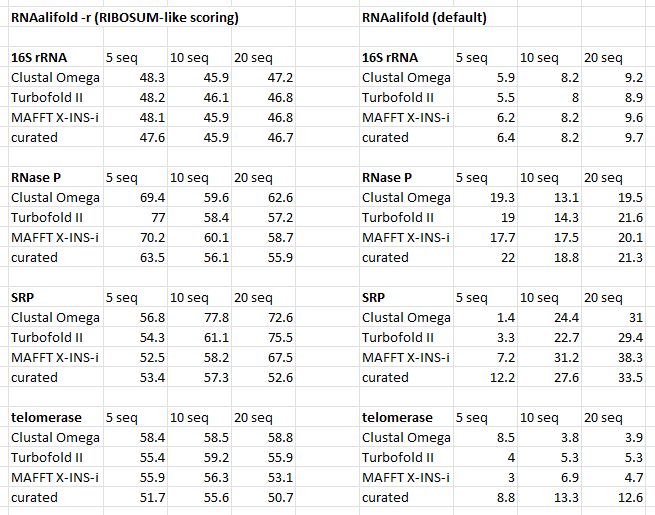

Table S13: Prediction accuracy of align-then-fold methods for RNase P and 16S rRNA. Tables are provided for curated and MAFFT X-INS-I alignments. Average PPV, sensitivity, and F1 scores are provided for 10 alignments consisting of 100 sequences each.

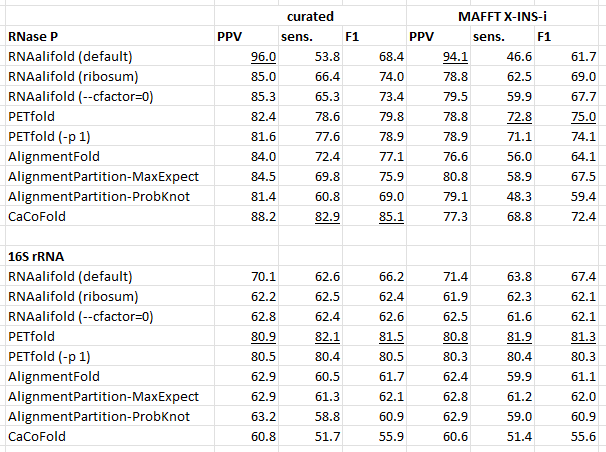

Table S14: AlignmentFold and AlignmentPartition average run time in seconds.

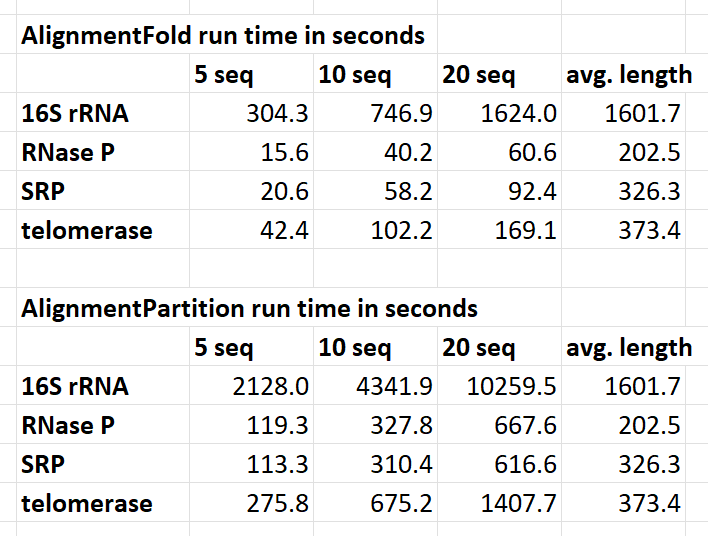

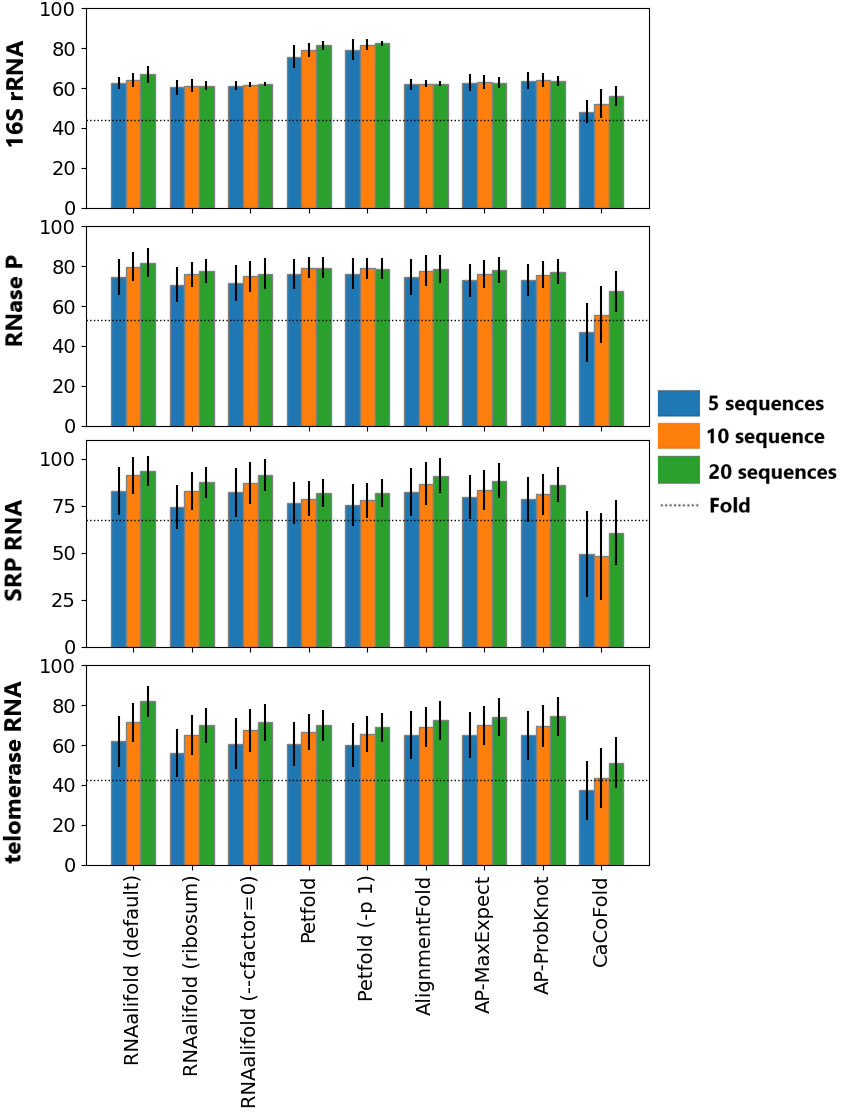

*Figure* *S1: Mean PPV for RNAalifold (default), RNAalifold (ribosum), RNAalifold (--cfactor=1), Petfold, Petfold (-p 1), AlignmentFold, AlignmentPartition-MaxExpect (AP-MaxExpect), AlignmentPartition-ProbKnot (AP-ProbKnot), and CaCoFold on 200 sets of 5, 10 and 20 sequences each aligned using MAFFT X-INS-i.* Plotted uncertainty is ± the standard deviation. *PPV of single sequence folding with RNAstructure Fold (free energy minimization) has been plotted using a dotted line.*

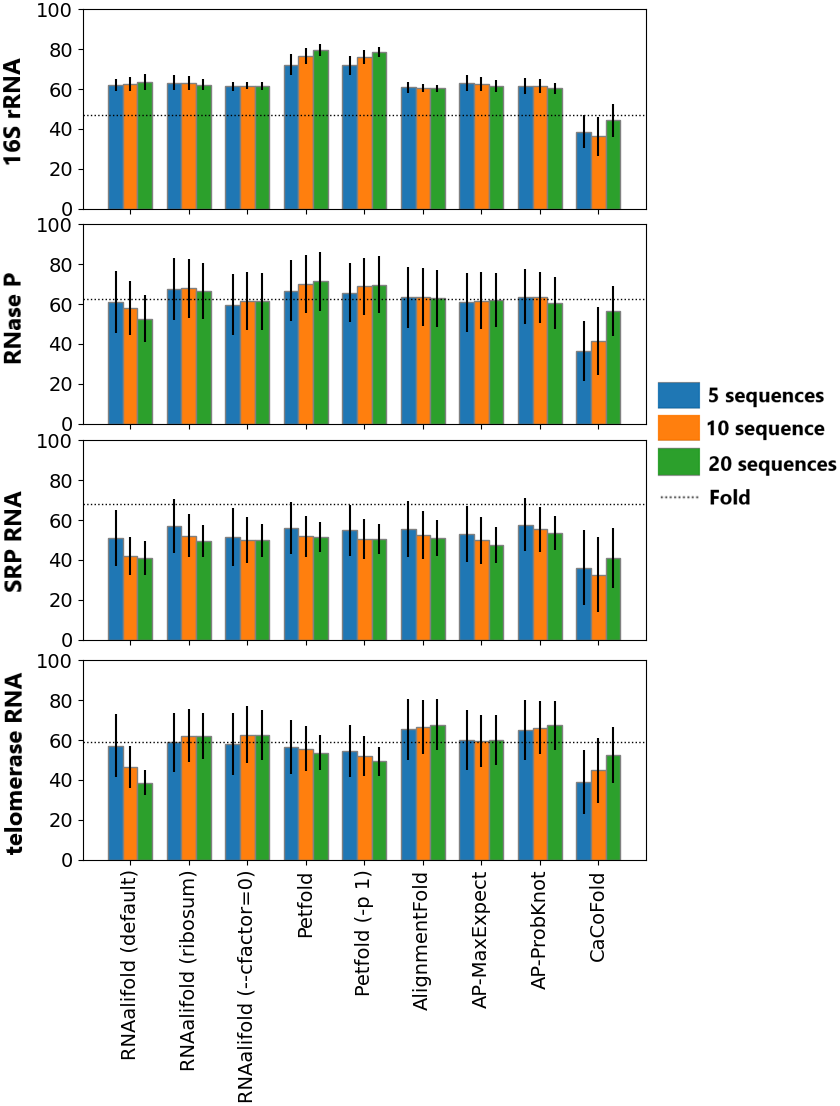

*Figure* *S2: Mean sensitivity for RNAalifold (default), RNAalifold (RIBOSUM), RNAalifold (--cfactor=1), Petfold, Petfold (-p 1), AlignmentFold, AlignmentPartition-MaxExpect (AP-MaxExpect), AlignmentPartition-ProbKnot (AP-ProbKnot), and CaCoFold on 200 sets of 5, 10 and 20 sequences each aligned using MAFFT X-INS-i.* Plotted uncertainty is ± the standard deviation. *Sensitivity of single sequence folding with RNAstructure Fold (free energy minimization) has been plotted using dotted line.*

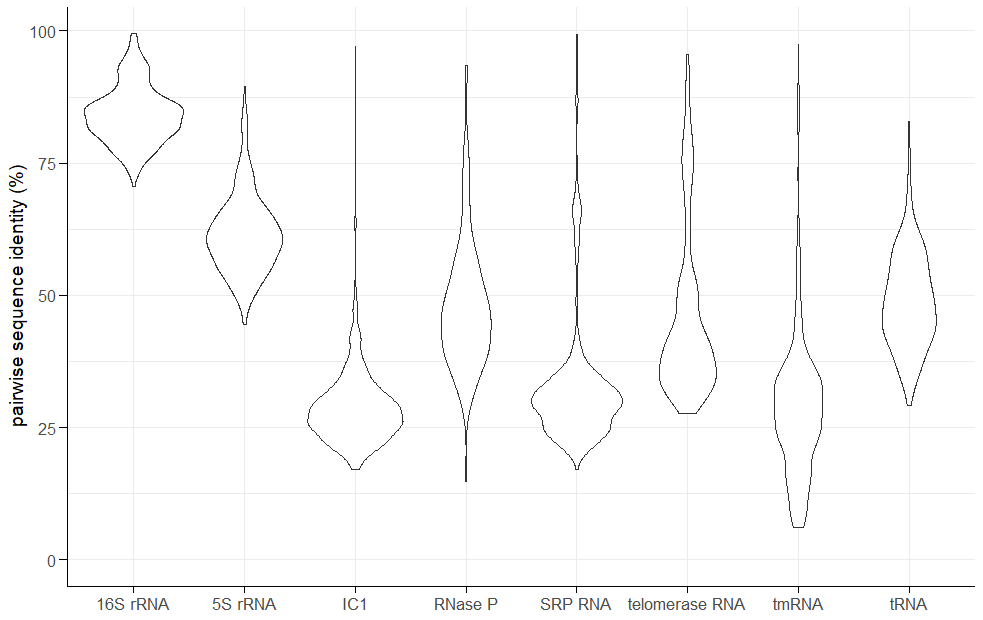

*Figure* *S3: Pairwise sequence identity for RNA families in RNAStrAlign dataset. 500 pairs of sequences were randomly sampled from curated alignments and percent sequence identity determined.*
